# Responses of bats to landscape transformation in an Andean agricultural landscape

**DOI:** 10.1101/555516

**Authors:** Mariana Vélez-Orozco, Laura M. Romero, John H. Castaño, Jaime A. Carranza-Quiceno, Jairo Pérez-Torres, Diego A. Torres

## Abstract

The current debate on the future of biodiversity gives rise to the need to integrate agricultural landscapes into conservation strategies. Bats are an important component of vertebrate diversity in many terrestrial landscapes where they provide invaluable and important ecosystem services to human societies such as insect pest control, pollination and seed dispersal. Here we study bat diversity and abundance in three landscapes representing a transformational gradient (continuous forests, forest fragments and crops) in an Andean agricultural scenario known as Colombia’s “Coffee Cultural Landscape”. We captured 1146 bats from 32 species and 4 families. The bat diversity and abundance in this landscape were high, especially for frugivorous bats, but there were no differences among the three transformational landscapes. However, some species were captured differentially between landscapes, suggesting that these landscapes have characteristics that influence the relative abundance of bats. Additionally, body weight and sex affect the abundance of some species in forest fragments and crops.

## Introduction

The current debate on the future of biodiversity in a context of accelerated species extinction rates (Ceballos et al. 2017) and ecosystem deterioration (Hansen et al. 2013), exacerbated by the insufficiency of protected areas for conservation (Chazdon et al. 2009), gives rise to the need to integrate agricultural landscapes into conservation strategies. These agricultural landscapes have generated a particular interest as reservoirs of biodiversity, since some of them may resemble natural ecosystems in terms of species diversity (Harvey et al. 2006) and in their capacity to generate ecosystem services (Landis 2017).

The structural and compositional heterogeneity of agricultural landscapes influences their capacity to maintain biodiversity (Fahrig et al. 2011). Simple landscapes, for instance, show a lower diversity of arthropods and vertebrates than more complex landscapes (Schmidt et al. 2005; Harvey et al. 2006; Frishkoff et al. 2014). Therefore, heterogeneous agricultural landscapes, in addition to maintaining productive activities, also favor the permanence of biodiversity and the supply of ecosystem services.

Bats are an important component of vertebrate diversity in many terrestrial landscapes. They provide important ecosystem services for human societies, such as insect pest control, pollination and seed dispersal (Kunz et al. 2011). The accelerated ecosystems transformations throughout the world, particularly because of deforestation, threaten many bat populations (Meyer et al. 2016) and the ecosystem services that they provide (García-Morales et al. 2016).

In the Andean region of Colombia, the country with the highest neotropical bat diversity known (Solari et al. 2013), more than 70% of the forest cover has been transformed to urban, agriculture or pasture (Etter and van Wyngaarden 2000). However, factors such as environmental variability and cultural diversity in the management of productive systems have generated a heterogeneous spatial mosaic where forest areas are mixed with cultivated areas, probably favoring the persistence of bat populations (Cardona et al. 2016).

The objective of this work was to study the bat diversity in a heterogeneous agricultural landscape in the Colombian Andes. Specifically we asked (1) if bat diversity is favored in this landscape and (2) how bats respond to three different transformational scenarios (continuous forests, forest fragments and crops). As bats are highly mobile, we expected no differences in overall bat diversity among transformational scenarios; however, since individual bats differ in functional traits (e.g. body weight, trophic guild and sex), we also expected species-specific responses to these transformative scenarios.

## Materials and methods

### Study area

The study area is located on the western slope of the Central Andes (Colombian Cordillera Central) of Colombia. UNESCO recognizes this region (including the departments of Riseralda, Quindío and Caldas) as a World Heritage site known as the “Colombian Coffee Cultural Landscape”, due to a particular combination of coffee-growing cultural practices and sustainability, example of outstanding human adaptation to difficult geographic conditions over which a homogenous mountainous coffee agriculture in a mixture of natural, economic and cultural elements have become highly and successfully blended in a sustainable manner. In this region, coffee has been produced for more than 100 years and is one of the most important crops in the area. Other important crops are pastures, bananas, vegetables, forest plantations and other fruits. The annual mean temperature oscillates between 16 - 24 °C, the mean relative humidity is 79% and the mean precipitation is 3358.4 mm.

We selected three landscapes representing a transformational gradient: (1) continuous forests, (2) forest fragments, which are immersed in a matrix of crops, and (3) crops without forests but with bamboo (*Guadua angustifolia*). Three sampling points were located in the center of a 1 km buffer of each landscape scenario totaling nine sampling localities. Each sampling point was separated by at least 4 km from other sampling points. All sampling points were located between an elevation of 1600 - 2000 meters.

### Bat sampling

Bats were captured between August 2016 and August 2017 using mist nets (12 meters long) located at 1 - 5 m above the ground. In total we accumulated 47012 net hours of sampling effort (forests: 9297; fragments: 19116; crops: 18599). Each captured bat was weighed, measured, sexed and marked with a consecutive number using tattoo pliers. Each bat was identified to species and a series of 41 individuals were collected as voucher specimens to be deposited in the mammalogy collection of the University of Santa Rosa de Cabal (Appendix I). Regional environmental authorities approved animal’s capture (CARDER- Corporación Autónoma Regional de Risaralda, license number 2004-Sep 2016) and the Animal Use and Care Committee of the University of Santa Rosa de Cabal approved all of the procedures.

### Data analyses

We used capture rate as an indication of bat abundance. Abundance was calculated as *CR=(i.n/m.h)*, where *i=*captured bats, *n=*sampling nights, *m=*number of nets, and *h=*sampling hours (Pérez-Torres and Ahumada 2004). To assess inventory completeness we divided the observed species richness between the species richness estimated by the index Jackknife 1, which was calculated with the software Estimates based on a matrix of species presence or absence and randomized 100 times (Colwell and Elsensohn 2014).

Capture rate was calculated for the three most abundant feeding guilds (nectarivorous, frugivorous and insectivorous) for species and for body weight intervals. We used a Chi-square to test differences in the capture of adult male or females using total abundances. Shannon-Wiener and Simpson indices were used to quantify the diversity between forests, fragments and crops. A Kruskal-Wallis test was used to find differences in capture rate and diversity among landscapes. Finally, the Jaccard index was used to explore the similarity between forests, fragments and crops. ANOVAs were run with the statistical software GraphPad Prism. The significance level was set at 5% (P<0.05). Data was expressed as mean ± standard deviation.

## Results

We captured 1146 bats from 32 species and 4 families (Table 1). Bats from five feeding guilds (frugivorous, nectarivorous, aerial insectivorous, gleaning animalivorous and hematophagous) were present; however, frugivorous bats were the richest and most abundant guild. The species richness estimator Jack 1 (Fig. 1A) suggested a greater richness than that observed, reaching a sampling completeness of 84%. The three landscapes showed a medium degree of similarity with forests and fragments being more similar to each other (66%) than to crops (60%) (Fig. 1B).

**Table 1.**
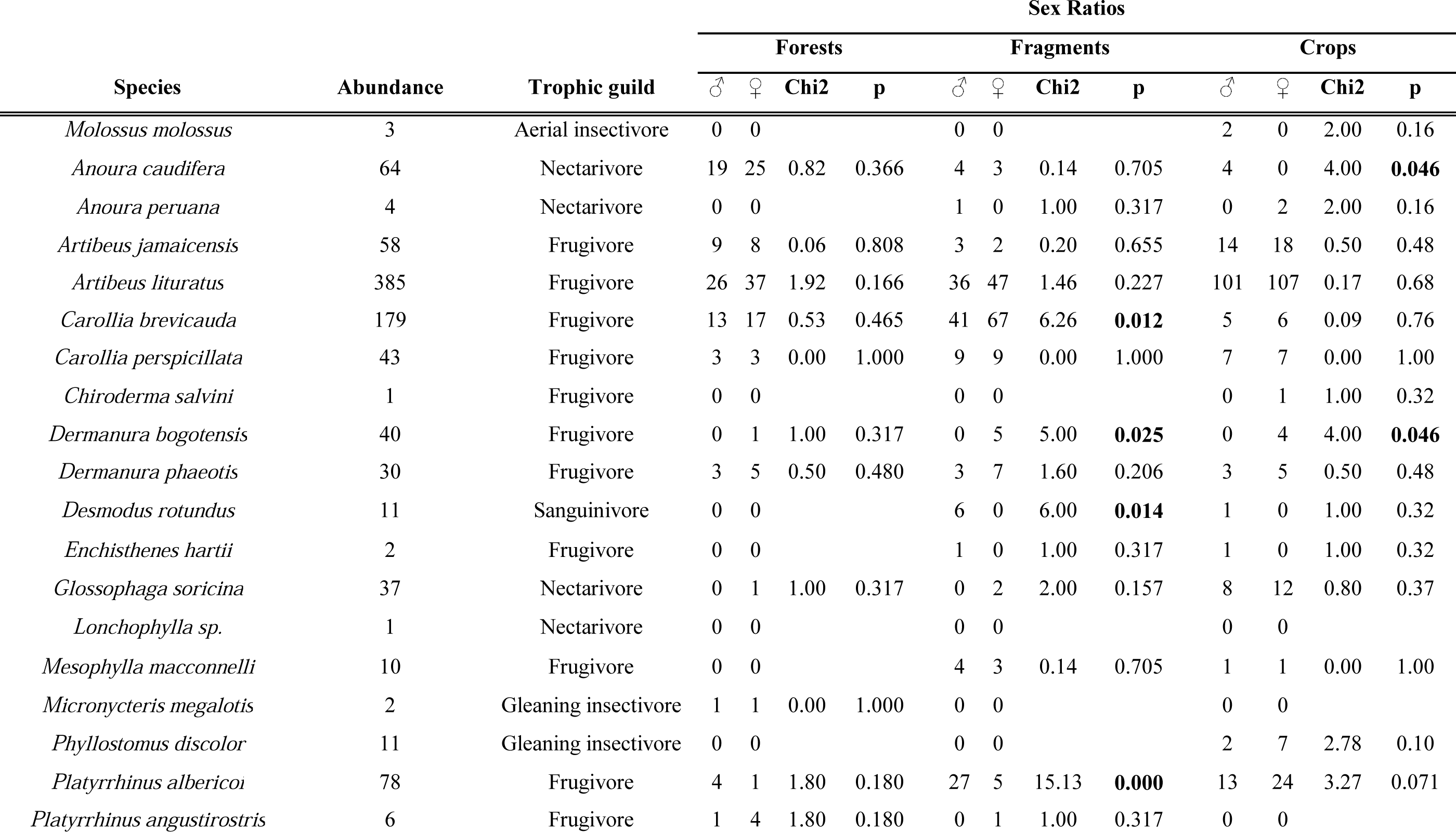
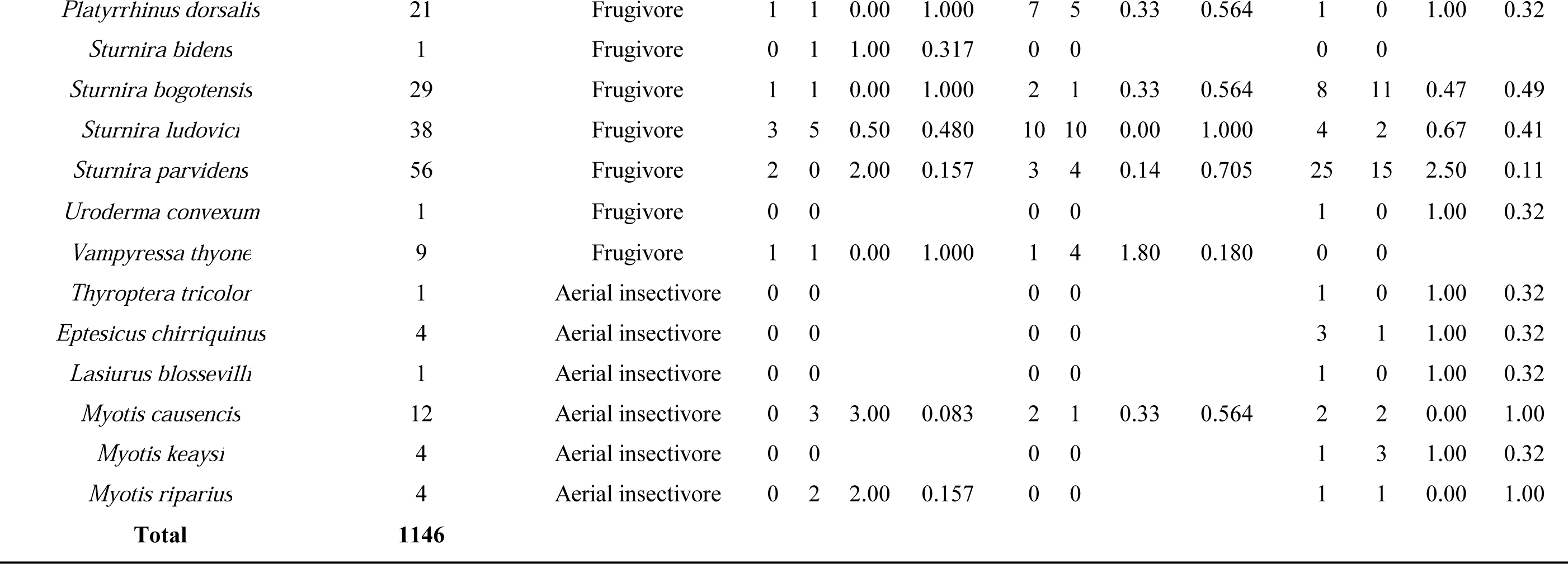
Species list, trophic guilds and sex ratios of captured bats in the Andean agricultural landscape. Sex ratio is based only in adult captures.

**Fig 1.**
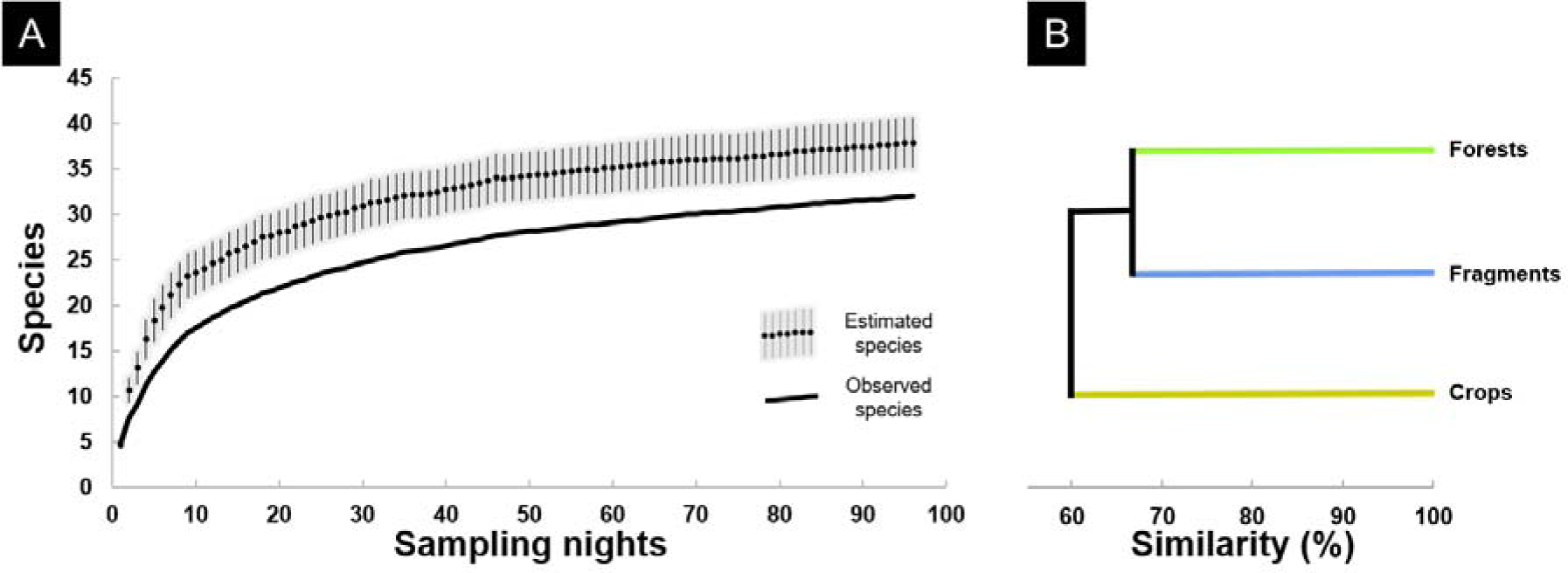
Curves of estimated (Jacknife 1 ± SD) and observed species richness (A). Dendrogram obtained with Jaccard similarity coefficient and average linkage method for the three landscape scenarios.

Neither the indices of diversity nor capture rates were different among the three landscapes (Fig. 2); however, there were differences when analyzed by species (Fig. 3A). *Anoura caudifera* was significantly more abundant in forests (H = 5.6; *P* = 0.04), *C. brevicauda* in fragments (H = 7.2; *P* = 0.003) and *S. parvidens* in crops (H = 5.96, *P* = 0.02). When pooled by feeding guilds, the capture rate of nectarivorous (Fig. 3B), frugivorous (Fig. 3C) and insectivorous bats (Fig. 3D) were the same among the three landscapes.

**Fig 2.**
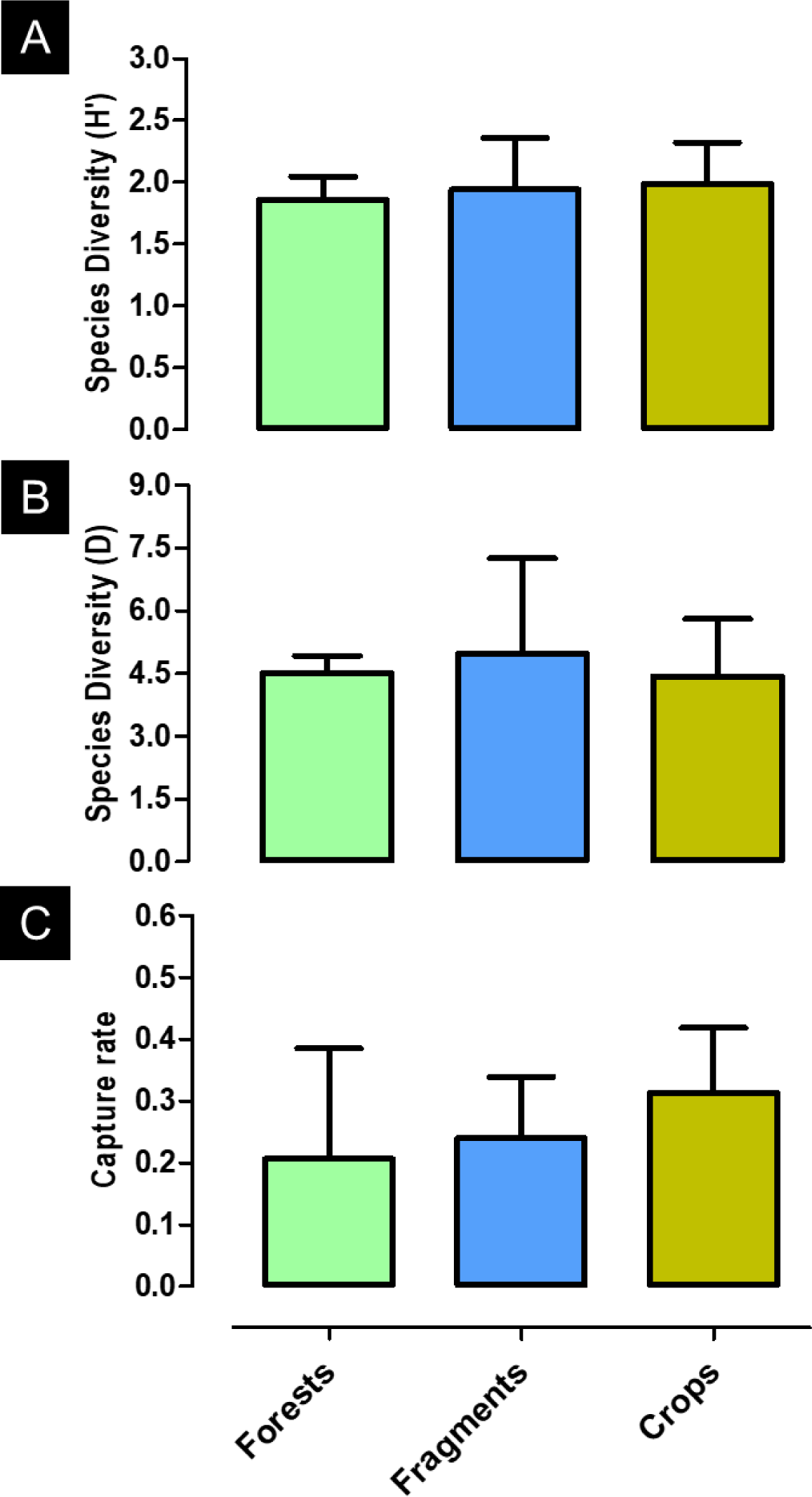
Shannon-Wiener (A) and Simpson (B) indices of diversity, and the relative abundance (capture rate) of all bat species (C) in the three landscape scenarios. Data are expressed as mean ± SD.

**Fig 3.**
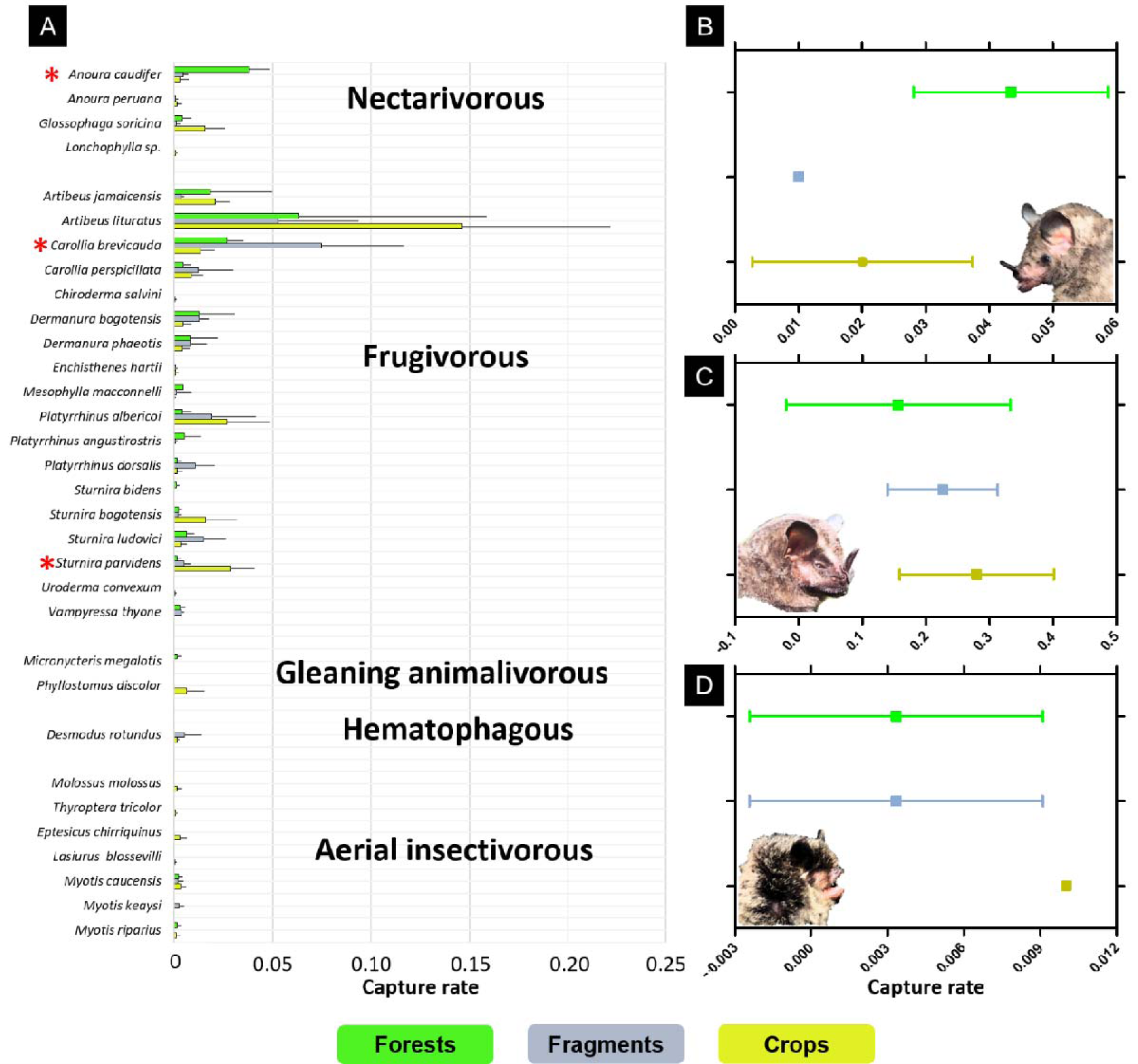
Relative abundance of species (captured rate) in each of the three landscape scenarios (A) and relative abundance pooled by the three most common trophic guilds: nectarivorous (B), frugivorous (C) and aerial insectivorous (D). Red asterisks indicate species with capture rates significantly different among the three landscape scenarios. Data are expressed as mean ± SD.

In four species the abundance of adult bats by sex varied between landscapes (Table 1). In fragments *C. brevicauda* and *D. bogotensis* females were more abundant than males and *P. albericoi* and *D. rotundus* males were more abundant than females. In crops *D. bogotensis* females and *A. caudifera* males were more abundant. In forests, differences in sex proportions were not detected. Finally, larger bats were captured more frequently in crops than in forests or fragments (H = 27.6, P = 0.0001) (Fig. 4A), particularly those with a body weight between 60 and 80 grams (H = 10.8, P = 0.02) (Fig. 4B).

**Fig 4.**
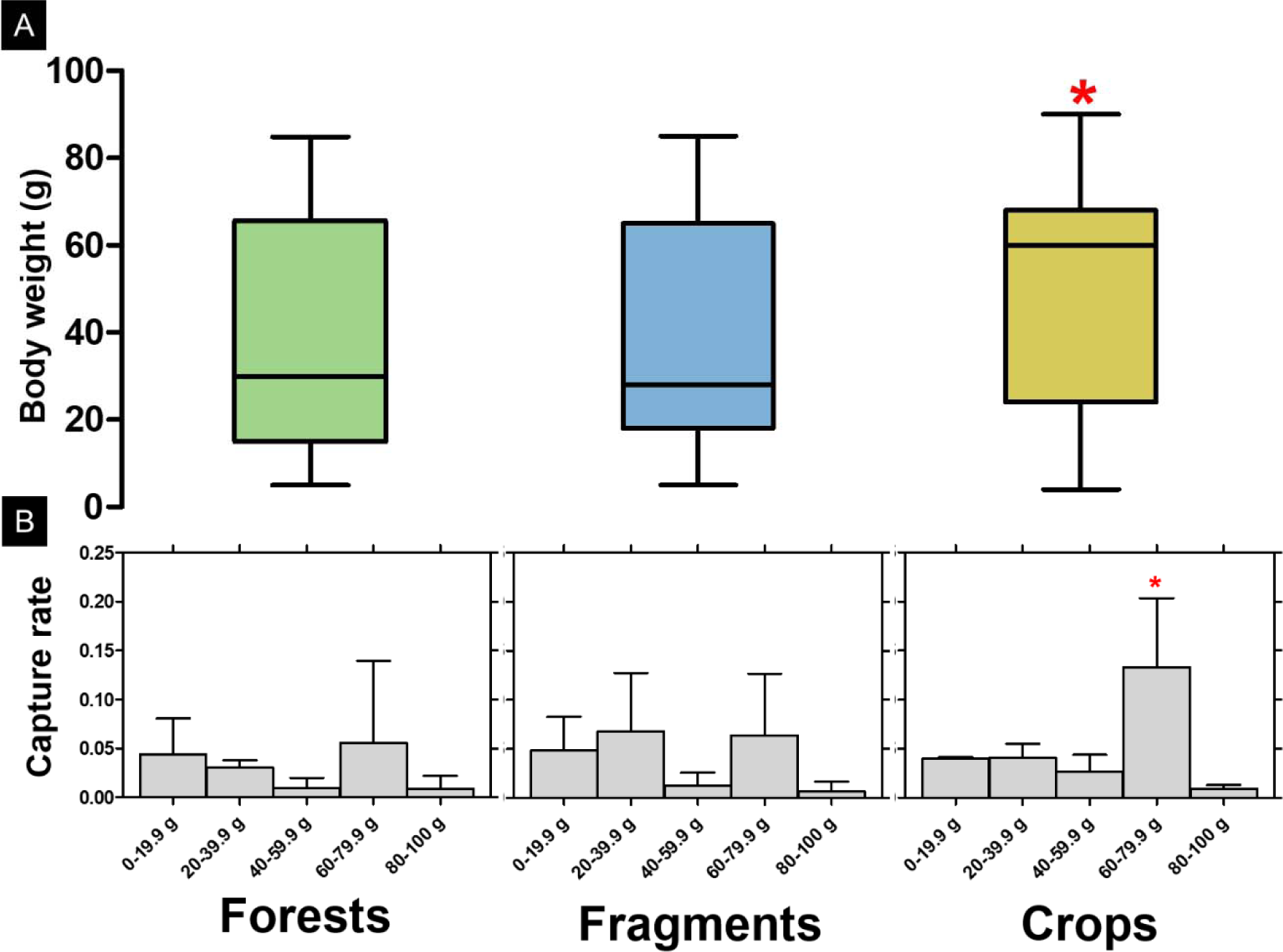
Body weight of all captured bats (A), excluding young and pregnant females, and the relative abundance (capture rate) of body weight intervals in each of the three landscapes (B). Red asterisks indicate species with capture rates significantly different among the three landscape scenarios. Data are expressed as mean ± SD.

## Discussion

The Colombian Coffee Cultural Landscape showed a high diversity of bats. As we predicted, there was no difference between continuous forests, forest fragments, and crops. This could be explained by the dispersal abilities of bats. However, some bat species were captured differentially between landscapes

The bat diversity found in this rural landscape is highly dominated by frugivorous bats. Bats were not only the most diverse in terms of species, but they were also the most abundant, which is the common pattern in tropical Andean forests (Soriano 2000) and coffee agricultural landscapes (Numa et al. 2005; Cardona et al. 2016). Furthermore, two frugivorous bats, *C. brevicauda* and *A. lituratus*, account for almost half of the captures. Aerial insectivorous bat diversity was probably underestimated because of the use of mist nets. We captured only six of the 16 aerial insectivorous bat species recorded for this area of the Colombian Andes (Castaño et al. 2018b), so relative abundances of this feeding guild between landscapes must be interpreted with caution.

Commuting flights of bats can reach several kilometers (Bernard and Fenton 2003), so it was early recognized that bats experience the landscape in a different way than other animals with less mobility (Moreno and Halffter 2000). For instance, individual bats might roost in forests, fly to crops and fragments to eat and return to the forests to roost (Bernard and Fenton 2003; Aguiar et al. 2014). This dispersal capacity could explain why no differences were found in diversity indices between scenarios of the landscapes studied.

*Artibeus lituratus*, with a body weight of 62.91±7.57 g (n=363) was the most captured species in crops. Two other large frugivorous bats, *P. albericoi* (62±7.35 g; n=61) and *A. jamaicensis* (65±8.64 g; n=41), were also frequently captured in this landscape. The presence of these three species explains why in crops the capture rate of bats having a body weight interval of 60-79.9 g was significantly high. The abundance of large bats in transformed ecosystems (e.g. urban areas) has been explained because of the availability of nutrients and by the abilities that larger bodies confer (Jara-Servín et al. 2017): for example stronger bites (Freeman and Lemen 2010) allow access to more food items. In our study landscape, for instance, *Cecropia angustifolia* is abundant in crops and other transformed areas. These trees fruit year-round (Zalamea et al. 2011) and are eagerly consumed by *Artibeus spp.* and *Platyrrhinus alberico* (Castaño et al. 2018a). Also, cultivated species such as guava (*Psidium guajava*) are abundant in crop areas, providing bats with many fruits.

Processes such as forest fragmentation or forest replacement by crops modify the vegetation composition and dynamics, favoring the recruitment and establishment of pioneer plants at the border of fragments (Zhu et al. 2004) and open areas such as *Solanum*, *Piper* and *Cecropia*. *Carollia brevicauda* and *S. parvidens* feed preferentially on the fruits of these plant species (Andrade et al. 2013; Castaño et al. 2018a), which might explain why these two species were captured more frequently in fragments and in crops. A similar relation to the composition of the vegetation might be drawn for the bat *A. caudifera*, captured mainly in forests. This is a nectarivorous species highly dependent on flowering plants that grow inside or at the border of forests (Muchhala and Jarrín-V 2002).

Differences in relative abundance of the sexes were detected in five species. This phenomenon has been found in many species of bats (Stoner 2001) and is generally attributed to foraging segregation and differences in habitat preferences; however, more data are needed to explore the causes of these patterns in our studied landscape.

Despite the establishment of fragmentation and crops, bat diversity in this Andean landscape remains high. Since not all bat species are favored by the same landscape scenarios, it is important to maintain diverse landscapes such as fragments, continuous forests, crops, gardens, pastures and forest plantations to favor the conservation of bats of different feeding guilds and the ecosystem services that they provide.

## Acknowledgments

We thank P Vélez, D.L. Rodríguez, C. Villabona, and C. López for their assistance in the field. We also thank the Corporación Autónoma Regional del Risaralda (CARDER), Central Hidroeléctica de Caldas (CHEC), landowners and farm administrators for research permission and collaboration. DAT thanks to COLCIENCIAS (Grant 775, Jóvenes Investigadores e Innovadores por la Paz del 2017).

## Funding

This work was supported by the Patrimonio Autónomo Fondo Nacional de Financiamiento para la Ciencia, la Tecnología y la Innovación Francisco José de Caldas [grant number 643771451270]; and the Vicerrectoría Académica Pontificia Universidad Javeriana [grant number 1215007010110].

## Conflict of interests

The authors declare that they have no conflict of interest.

